# CyFj11: FlowJo v11 Workspace Import and Legacy Format Export for R-Based Flow Cytometry Analysis

**DOI:** 10.64898/2026.07.28.741192

**Authors:** Bernd Jagla, Slobodan Culina, Francois Le Guerroue, Esma Karkeni, Milena Hasan

## Abstract

High-dimensional flow cytometry measures immune cells at single-cell resolution, enabling systematic characterization of cell populations at scale. But harnessing this potential requires seamless interoperability between the interactive gating tools used by biologists and the statistical environments used for detailed downstream analysis. FlowJo, one of the most widely used commercial cytometry analysis software packages, now stores workspaces in a format that existing R tools cannot read, leaving researchers unable to import their gating strategies into R, or to return R-based results to FlowJo for visual review or collaborative sharing, without manual reconstruction. We present CyFj11, an R package that closes this gap, enabling import of FlowJo v11 gating hierarchies into R and export of R-defined gates back to FlowJo (throughout this paper, “import” refers to bringing a FlowJo v11 workspace into R, and “export” to writing an R-derived GatingSet back out to FlowJo). Using a combination of synthetic test scenarios and a real-world immunophenotyping dataset, we show that population counts in FlowJo 10 and 11 matched R-derived values with Pearson correlation coefficients exceeding 0.99. CyFj11 is platform-independent, requires no additional software infrastructure, and is freely available at https://github.com/C3BI-pasteur-fr/CyFj11.

## Introduction

Flow cytometry is a cornerstone technology for characterizing immune cell populations at single-cell resolution. As datasets grow in dimensionality and cohort sizes scale to hundreds of donors, the ability to apply systematic, reproducible gating strategies across samples, and to integrate gating results with statistical and genomic analyses, becomes essential. Population-scale studies linking immune cell phenotypes to genetic variation (1) exemplify both the scientific opportunity and the interoperability barrier such workflows face: gating defined in interactive tools has to reach the statistical environments where phenotypes are linked to genetic data, a step CyFj11 is built to enable; without it, count tables have to be manually exported.

FlowJo (2) is one of the most widely used interactive commercial software packages for flow cytometry data analysis. Its version 11, released in 2025, adopts a JSON-based workspace format that no current R/Bioconductor tool can read. CytoML v2.24.0 (3), the primary Bioconductor gateway for FlowJo data, reads only the older v10 XML format, and its FlowJo export function (gatingset_to_flowjo()) depends on an external Docker, which is currently not compatible with Apple Silicon Macs. Researchers who gate in FlowJo v11 therefore lack a reliable pathway to import gating hierarchies into R or to return R-derived results to FlowJo without manual reconstruction. Import matters just as much: bringing FlowJo v11 gating hierarchies into R lets the same gating strategy be applied systematically across large cohorts and fed directly into R/Bioconductor’s statistical and genomic analysis tools, rather than requiring each sample’s gates to be manually re-derived. Export from R into FlowJo is important since returning R-defined gates to FlowJo enables biologists to visually inspect and quality-control gates produced by automated or code-based pipelines, to share reproducible gating strategies with collaborators who work primarily in FlowJo, and to reconcile scripted analyses with the interactive review that remains central to cytometry practice.

CyFj11 addresses this with two functions: fj11_to_gatingset(), which imports a v11 workspace as GatingSet objects (4), and export_flowjo10_workspace(), which writes a GatingSet to FlowJo v10-compatible XML (Figure 1). The new, undocumented JSON format relies on a complex web of cross-referenced identifiers across multiple components, making faithful generation from R prohibitively difficult. Rather than attempting to write native FlowJo v11 files, CyFj11 exports to the well-documented v10 XML format and relies on FlowJo v11’s built-in ability to open v10 workspaces, a capability that FlowJo v11 supports natively. Full implementation details, including the FlowJo v10/v11 format comparison, gate-type conversion, and transformation parameter mapping, are provided in Appendix S4. The package requires no external dependencies beyond standard Bioconductor infrastructure, is fully compatible with openCyto v2.22.0 (5) and flowWorkspace v4.22.1.

**Figure 1.**
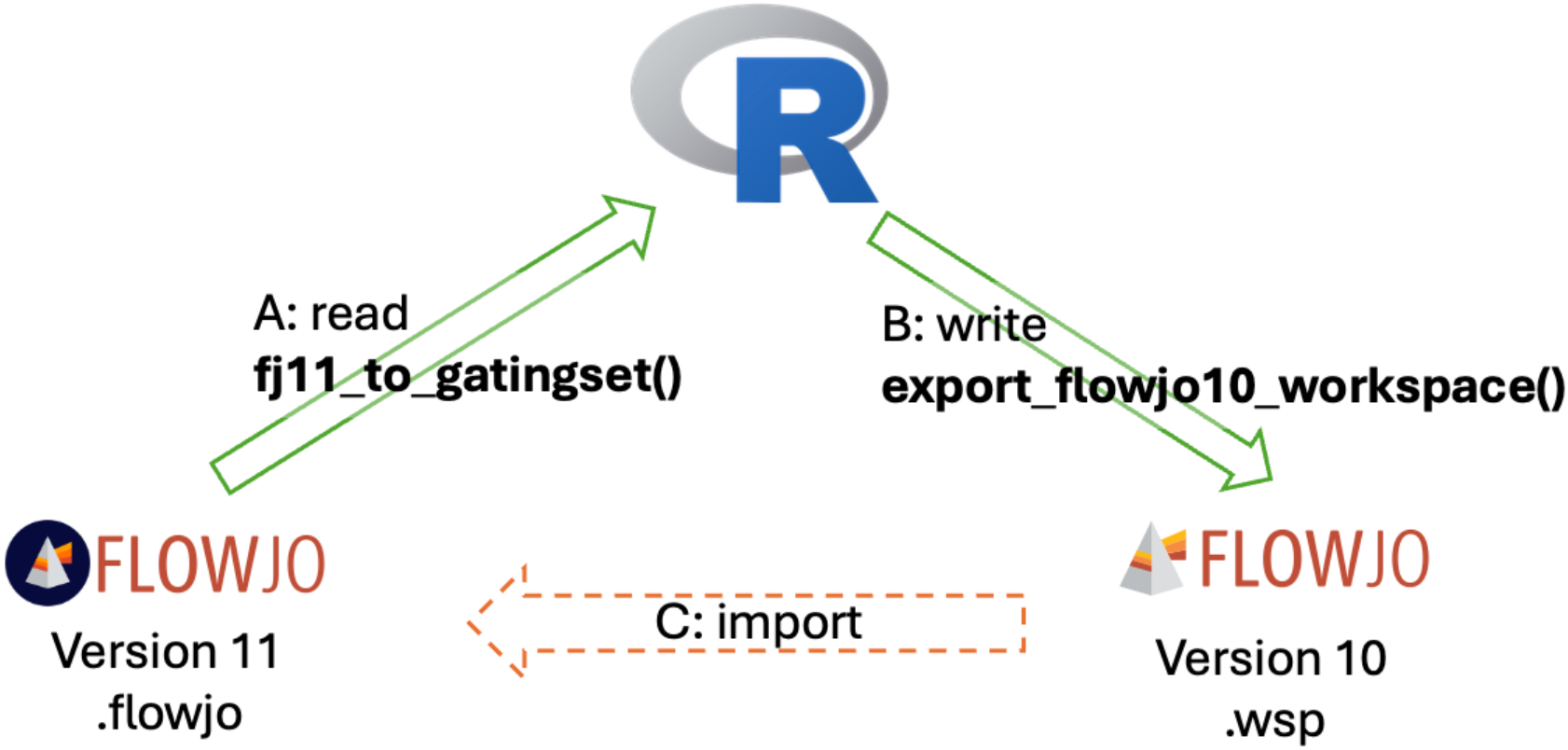
CyFj11 workflow overview. Arrow A: fj11_to_gatingset() reads a FlowJo v11 .flowjo workspace into R as a GatingSet object. Arrow B: export_flowjo10_workspace() writes a GatingSet to a FlowJo v10-compatible .wsp XML file. Arrow C (dashed): the exported .wsp can be opened in FlowJo v11; this step is performed outside R and is subject to FlowJo v11’s partial backward compatibility with the v10 format (see Limitations).

## Methods

**read_flowjo11_workspace**() extracts and parses the .flowjo file, which is a ZIP archive containing a JSON workspace. It reconstructs the workspace components — sample groups, FCS file references, gate definitions, per-sample gate variations, and transformation parameters — and returns them as a structured list for downstream processing.

**fj11_to_gatingset**() takes the parsed workspace and constructs flowWorkspace GatingSet objects. It locates FCS files by recursive directory search, applies compensation, converts transformation parameters to flowCore objects, and imports the full gate hierarchy. Gate coordinates are converted from FlowJo’s display space to the raw data space expected by flowCore. Because FlowJo v11 supports per-sample tailored gates, the function returns a list of per-sample GatingSet objects; users who require a single merged object can call flowWorkspace::merge_list_to_gs().

**export_flowjo10_workspace**() serializes a GatingSet to FlowJo v10 XML, replicating FlowJo v10’s native element structure, namespace conventions, and transformation encodings. Supported gate types include rectangle, polygon, ellipsoid, range, and boolean. Compensation matrices and all transformation parameters are preserved. Per-sample gate variations cannot be represented in v10 XML and are respectively omitted or collapsed to group-level on export (see Limitations).

## Results

CyFj11 was validated across the full round trip it is designed to support: R-defined gates were exported to a FlowJo v10 workspace, that workspace was opened and re-saved in FlowJo v11, and the resulting v11 workspace was re-imported into R. At each stage, population counts were compared against the counts recorded in the reference program, providing a direct measure of how faithfully gate geometry and the associated cell populations are preserved across formats. The sections below report, in turn, export to FlowJo v10, a real-world dataset round trip (Test-14), import back into R from FlowJo v11, and Gating-ML standards compliance.

### Validation design

Thirteen synthetic test scenarios were constructed to systematically validate all supported gate types (rectangle, polygon, ellipsoid, range, boolean) and transformation types (linear, biexponential, log, arcsinh) in isolation and combination, including hierarchical and boolean configurations (full descriptions in Appendix S1). For each scenario, synthetic FCS files were generated in R, a GatingSet was constructed, and the result was exported to a FlowJo v10 XML workspace using export_flowjo10_workspace(). Each exported workspace was then opened in FlowJo v10 to manually record population counts for comparison against the R ground truth. The workspace was subsequently re-saved from within FlowJo v10; both the original CyFj11 exported version and the FlowJo v10 saved version were loaded with FlowJo v11, ensuring that any import failures observed in FlowJo v11 could be attributed to FlowJo v11’s handling of v10-format workspaces in general, rather than to any idiosyncrasy of the CyFj11-generated XML specifically; no major discrepancies were observed between the two versions once loaded into FlowJo v11. To assess FlowJo v11 compatibility, this workspace was imported into FlowJo v11, manually verified, and corrected where necessary (see Appendix S2 for details). The corrected workspace was saved in v11 format, and population counts recorded within FlowJo v11 at this stage served as the ground truth for the import-to-R comparison. This v11 workspace was then imported into R using read_flowjo11_workspace() and fj11_to_gatingset(), and the resulting GatingSet counts were compared against those FlowJo v11 reference counts. An additional test (Test-14) used the FlowSOM (6) example dataset — a mouse bronchoalveolar lavage (BAL) sample stained with a T/B/NKT/NK lymphocyte panel (15 populations across scatter and compensated fluorescence channels) to assess performance on a real-world dataset and to validate compensation matrix preservation through the round trip. This example dataset ships with the FlowSOM package (6); we selected it because bundling with a widely used Bioconductor package lets readers reproduce the test without additional downloads, and its lineage-marker panel (including NK1.1, CD45, TCRβ, and TCRγδ) exercises both gate-geometry fidelity and compensation handling across a biologically meaningful gating hierarchy. All source code, FCS files, and workspaces are deposited at https://doi.org/10.5281/zenodo.20933199. The package is maintained at https://github.com/C3BI-pasteur-fr/CyFj11, pending acceptance into Bioconductor.

### FlowJo v10 export

For all gate types (rectangle, polygon, boolean, and ellipse), FlowJo v10 reproduced R-derived counts with high fidelity (Pearson correlation coefficients > 0.99, Figure 2) across all tests. Rectangle and range gates showed exact agreement in every case. The small count differences that did appear (at most 14 cells, or 0.2%, confined to polygon and ellipse gates) arise because cells that fall very close to a gate boundary line may be counted as inside or outside depending on how each program handles that exact boundary value; the gate itself is in the right place. This was confirmed by extracting gate coordinates directly from the exported file: boundary coordinates matched the R definitions exactly across all axis transformation types tested. To put the 14-cell maximum in perspective: a biologist redrawing the same gate by hand on two separate occasions would not place it identically and could easily shift at least as many cells (7).

**Figure 2.**
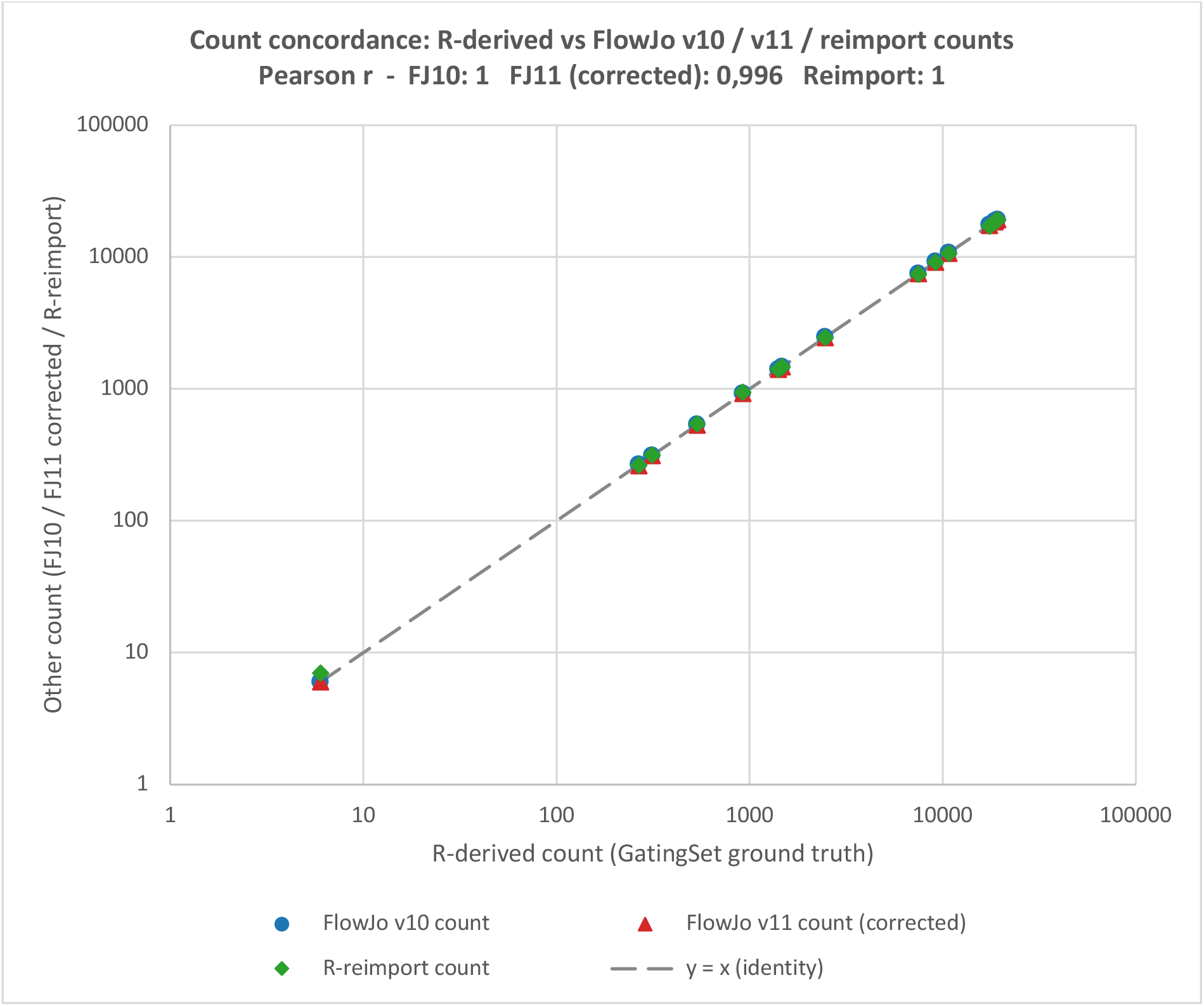
Count-concordance between R-derived and FlowJo-derived population counts. Scatter plot of population counts per test (R/CyFj11-derived count vs. FlowJo count); the diagonal denotes the line of identity. Points close to the line indicate high agreement between programs.

In Test-14, all 12 populations on compensated fluorescence channels, including NK cells, B cells, T cell subsets, and CD4/CD8 populations, were reproduced with identical counts. The three scatter-channel populations differed by at most 10 cells (0.05%). The exact reproduction of fluorescence-channel counts is itself evidence that the compensation matrix was correctly transferred: obtaining the same cell counts after compensation in two different programs requires that both use the same spillover matrix and apply it with equivalent compensation arithmetic; residual rounding or floating-point differences between R and FlowJo (or between operating systems) could in principle shift borderline cells, but none was observed on these channels. This was confirmed directly by comparing the exported compensation matrix entry by entry against the original.

### FlowJo v11 import

Two systematic import failures restricted the populations available for count comparison. Some boolean gates and gates on arcsinh-transformed axes were silently dropped or incorrectly rendered when FlowJo v11 read the v10 workspace files; the same file opened without error in FlowJo v10, confirming the export is structurally correct. For rectangle and range gates on linear, log, and biexponential transformations, R-derived counts reliably reproduced the FlowJo v11 counts. Where discrepancies appeared, they were confined to polygon and ellipse gates, reflecting differences in how FlowJo v11 renders these shapes when reading v10-format coordinates rather than errors in the gate definitions themselves. For the real-world Test-14 dataset, FlowJo v11 was unable to open the original FlowSOM-distributed workspace file correctly; the same data opened correctly once exported via export_flowjo10_workspace(), indicating that CyFj11’s v10-XML output is more broadly compatible with FlowJo v11 than the original file format. This is consistent with a compensated-channel naming requirement identified: FlowJo v11 expects compensated $PnN keywords to carry a doubled “Comp-Comp-” prefix, which CyFj11’s export supplies but which the original, unmodified file does not (Appendix S4). We then compared to what FlowJo v11 read by reading that .flowjo (JSON) file using CyFj11 and exported as FlowJo v10 and reopening in FlowJo v10. FlowJo v10 reproduced R-derived counts to within 10 cells (≤0.05%) across all 15 populations, whereas FlowJo v11 still disagreed by up to 107 cells (0.58%), likely reflecting FlowJo v11’s own interpretation, rounding, or recomputation of the compensation matrix, distinct from the naming-related dropout affecting the twelve compensated-fluorescence populations. Figure 3 illustrates this directly for the Singlets2 gate, the population with the largest FJ11 divergence. Panel (a) shows the gate as rendered by FlowJo v11 (after importing the CyFj11-exported v10 workspace), the reference point for this FJ11-import comparison. Panel (b) shows the corresponding GatingSet gate in R after read_flowjo11_workspace() re-imported that FlowJo v11-saved file, representing what FlowJo v11 had actually understood (see figure3_singlets2_panels.R for source code to generate figure b). Panel (c) shows the same gate after this GatingSet was exported once more via export_flowjo10_workspace() and read by FlowJo v10. Although the three panels are visually almost identical, the underlying counts are not identical: for this gate, FlowJo v11 (a) differs from both the R/CyFj11 GatingSet (b) and FlowJo v10 (c) by several tens of cells, whereas (b) and (c) differ from each other by only a handful of cells. The same pattern holds across most gates in this dataset (Appendix S2, B.4.6), indicating that the residual divergence originates specifically in FlowJo v11’s own handling of the workspace rather than in the CyFj11 export/import pipeline.

**Figure 3.**
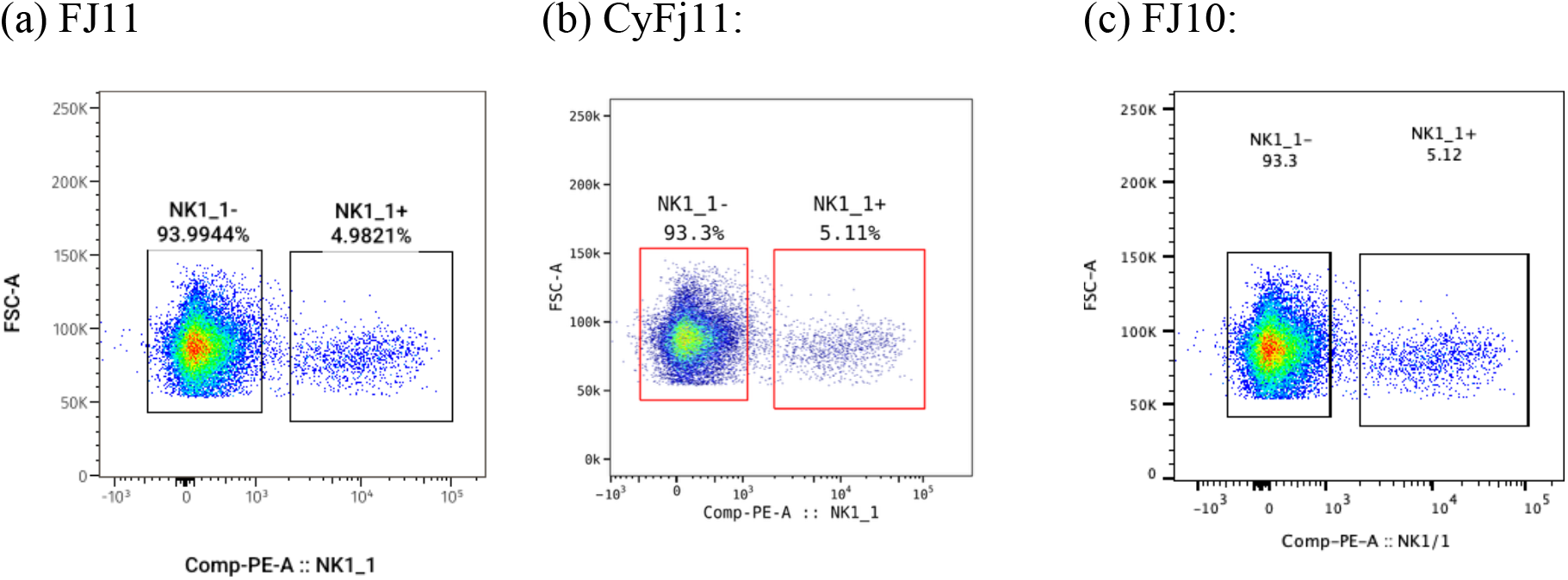
Visual comparison of the critical gate showing the largest FlowJo v11 discrepancy in Test-14 (the Singlets2 gate, differing by up to 107 cells / 0.58% from the R-derived reference), traced through a further re-import/re-export cycle. (a) The gate as rendered by FlowJo v11 after importing and saving the CyFj11-exported v10 workspace. (b) The corresponding gate on the GatingSet object in R, obtained by re-importing that FlowJo v11-saved workspace with read_flowjo11_workspace(). (c) The same gate as rendered by FlowJo v10 after this GatingSet was exported again via export_flowjo10_workspace().

**Table 1.**
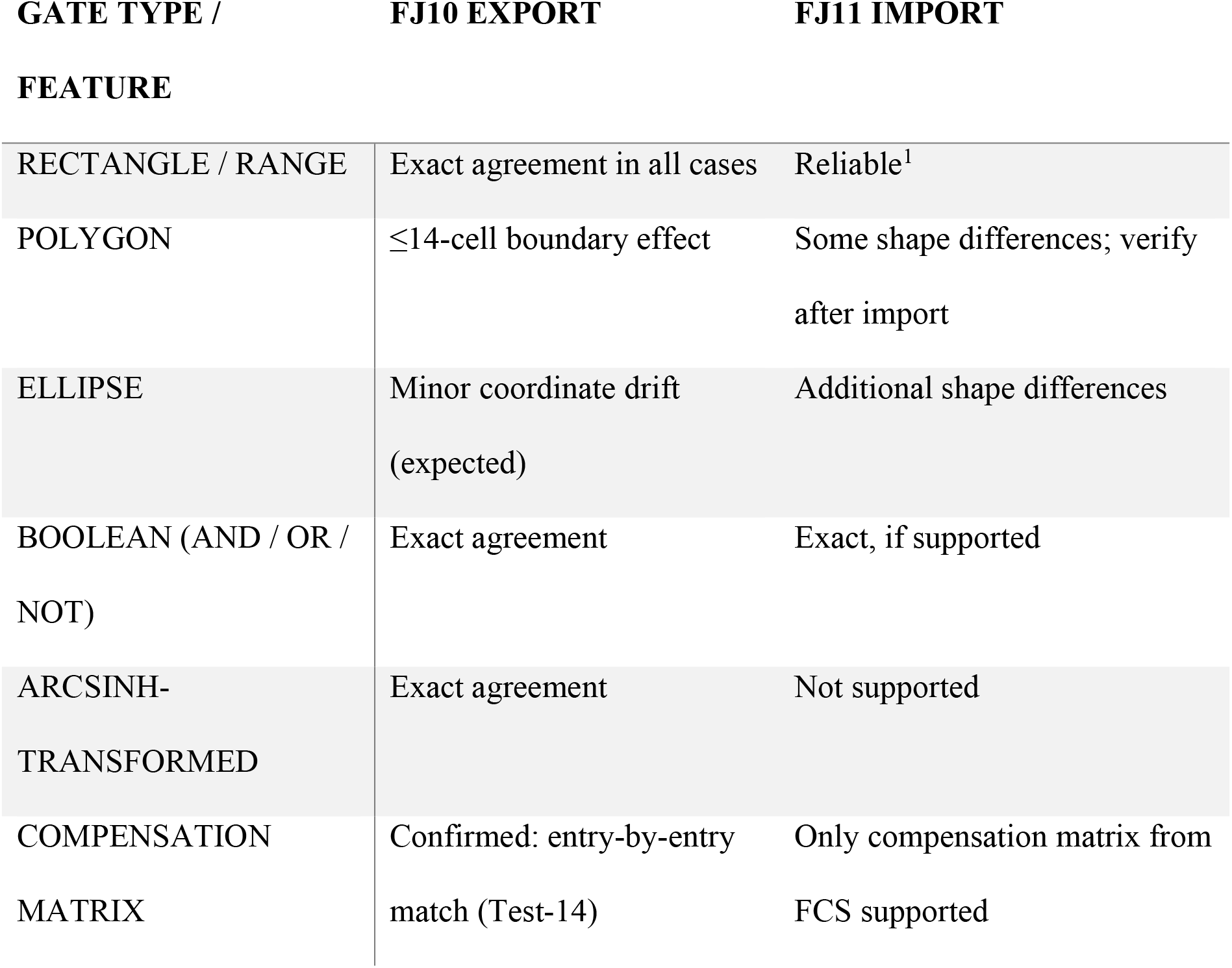
Export validation summary by gate type and feature.

### Standards compliance

All 50 applicable Gating-ML 2.0 structural checks passed across the 13 test workspaces (Appendix S3).

## Discussion

### Comparison with existing approaches

To our knowledge, CyFj11 is the first tool to import FlowJo v11 workspaces into R; because no current R/Bioconductor package can parse the v11 JSON format, a direct head-to-head comparison is not possible for the import direction, and correctness was instead assessed against FlowJo’s own population counts as ground truth. For export, the closest existing solution is CytoML’s gatingset_to_flowjo(), which produces FlowJo-compatible workspaces but depends on an external Docker image rather than running natively in R. CyFj11 removes this dependency, running entirely within R on standard Bioconductor infrastructure, and its v10-format output is designed to open directly in FlowJo v11 through that program’s backward compatibility, closing the round trip. Because both routes ultimately yield v10-format XML that is verified inside FlowJo, FlowJo’s native counts, rather than another converter’s output, are the appropriate reference for measuring fidelity, which is the basis of the validation reported above.

Beyond closing this specific import barrier, CyFj11 provides a complete, dependency-free round trip between R and FlowJo. fj11_to_gatingset() reconstructs full gating hierarchies, including per-sample tailored gates, all standard gate geometries, and transformation parameters, as native GatingSet objects ready for downstream analysis in R/Bioconductor. export_flowjo10_workspace() reverses this, writing GatingSets back to well-documented, Gating-ML 2.0–compliant v10 XML that opens natively in both FlowJo v10 and, through its backward compatibility, FlowJo v11, letting biologists visually inspect and quality-control gates produced by automated or code-based pipelines. Because the package runs entirely within R on standard Bioconductor infrastructure, it requires no external services, and the validation reported above shows this round trip preserves population counts, gate geometry, and compensation matrices with high fidelity.

Four limitations apply.

1. FlowJo v11’s v10 importer sometimes drops Boolean gates (observed in hierarchical Tests 11–13, not in the root-level Test-10) and mis-renders arcsinh ellipsoid gate boundaries (Test-13 FITC_PE_gate). In every case the exported v10 XML itself was reproduced correctly by FlowJo v10, localizing the visible failures to FJ11’s v10 importer. These incompatibilities likely affect any v10-format workspace opened in FJ11, not only CyFj11 exports.
2. Compensation matrix export has been validated indirectly and by element-wise comparison for Test-14. Manually modified compensations are not supported at the moment in CyFj11. In general, because of the complexity of the JSON format, it cannot be assured that all features are currently supported. Here we tried to cover the most common use cases.
3. Per-sample tailored gates are collapsed to group-level on v10 export.
4. Quadrant gates are not currently supported in the export functionality.

The residual differences observed across the validation pipeline underscore an inherent fragility in flow cytometry analysis more generally. Whether populations are gated manually in a GUI or defined automatically in code, the final counts are shaped by numerical precision, transform conventions, and file-format idiosyncrasies. Thus, even fully reproducible, automated workflows should be expected to yield small, non-zero discrepancies when compared across software platforms. FlowJo v11 remains under active development, with new releases roughly every 3–5 months, and several of the FJ11-specific import limitations noted above may be resolved in future versions.

## Supporting information

Appendix S1. Validation test scenarios: synthetic FCS data generation, transformation defaults, the fourteen-test scenario summary, source files and l

Per-population validation results, including Table S1 (per-population counts for FlowJo v10 export and FlowJo v11 import across all test scenarios)

Gating-ML 2.0 standards-compliance results: the 50 applicable structural checks across the 13 test workspaces

Technical implementation details: FlowJo v10/v11 workspace format comparison, gate-type conversion, transformation parameter mapping, and package arch

Package reference manual, generated from the package's roxygen2 documentation

## Acknowledgments

We thank Lorenzo Bonaguro for his careful reading of the manuscript

## Availability

CyFj11 is available as open-source software at https://github.com/C3BI-pasteur-fr/CyFj11. Validation datasets and scripts are deposited at https://doi.org/10.5281/zenodo.20933199. The package is compatible with R ≥ 4.0 and Bioconductor ≥ 3.14. It can be installed directly from the repository in R with remotes::install_github(“C3BI-pasteur-fr/CyFj11”). Installation instructions, a worked example using the bundled dataset, and full function documentation are provided in the package README and vignette. A static reference manual is also provided as Appendix S5. All results reported here were produced with CyFj11 v0.99.95 on R 4.5.0 and Bioconductor 3.22 (flowCore 2.22.1, flowWorkspace 4.22.1, openCyto 2.22.0, CytoML 2.24.0, and FlowSOM 2.18.0) under macOS 26.5.2 on Apple Silicon (M3/M4).

## Supporting Information

Additional supporting information may be found in the online version of this article.

**Appendix S1**. Validation test scenarios: synthetic FCS data generation, transformation defaults, the fourteen-test scenario summary, source files and locations, validation methodology and acceptance criteria, and regeneration instructions (Microsoft Word, .docx).

**Appendix S2**. Per-population validation results, including Table S1 (per-population counts for FlowJo v10 export and FlowJo v11 import across all test scenarios) (Microsoft Word, .docx).

**Appendix S3**. Gating-ML 2.0 standards-compliance results: the 50 applicable structural checks across the 13 test workspaces (Microsoft Word, .docx).

**Appendix S4**. Technical implementation details: FlowJo v10/v11 workspace format comparison, gate-type conversion, transformation parameter mapping, and package architecture (Microsoft Word, .docx).

**Appendix S5**. Package reference manual, generated from the package’s roxygen2 documentation (Microsoft Word/PDF).

**Data S1**. Source code, synthetic and real-world FCS files, and FlowJo v10/v11 workspaces used for the reported validation, archived at https://doi.org/10.5281/zenodo.20933199 and produced with the software versions listed under Availability.

## Footnotes

1 Visually indistinguishable from the R- and FlowJo v10-rendered versions of the same gate; on real-world data (Test-14) these gates nonetheless show measurable count differences (up to ∼0.6%; see Figure 3), likely reflecting FlowJo v11’s own recomputation of the compensation matrix rather than gate-geometry error. Treated here as expected inter-program variation rather than a reliability failure.

## References

1. The Milieu Intérieur Consortium, Patin E, Hasan M, Bergstedt J, Rouilly V, Libri V, et al. Natural variation in the parameters of innate immune cells is preferentially driven by genetic factors. Nat Immunol. 2018 Mar;19(3):302–14. doi:10.1038/s41590-018-0049-7

2. FlowJo TM, version 10 & 11. Becton, Dickinson and Company.

3. Finak G, Jiang W, Gottardo R. CytoML for cross-platform cytometry data sharing. Cytometry Part A. 2018;93(12):1189–96. doi:10.1002/cyto.a.23663

4. Greg Finak MJ. flowWorkspace [Internet]. Bioconductor; 2017 [cited 2026 Jul 6]. Available from: https://bioconductor.org/packages/flowWorkspace doi:10.18129/B9.BIOC.FLOWWORKSPACE

5. Finak G, Frelinger J, Jiang W, Newell EW, Ramey J, Davis MM, et al. OpenCyto: An Open Source Infrastructure for Scalable, Robust, Reproducible, and Automated, End-to-End Flow Cytometry Data Analysis. Prlic A, editor. PLoS Comput Biol. 2014 Aug 28;10(8):e1003806. doi:10.1371/journal.pcbi.1003806

6. Van Gassen S, Callebaut B, Van Helden MJ, Lambrecht BN, Demeester P, Dhaene T, et al. FlowSOM: Using self-organizing maps for visualization and interpretation of cytometry data. Cytometry Part A. 2015 Jul;87(7):636–45. doi:10.1002/cyto.a.22625

7. Finak G, Langweiler M, Jaimes M, Malek M, Taghiyar J, Korin Y, et al. Standardizing Flow Cytometry Immunophenotyping Analysis from the Human ImmunoPhenotyping Consortium. Sci Rep. 2016 Feb 10;6(1):20686. doi:10.1038/srep20686

