## Appendix S1. Validation test scenarios: synthetic FCS data generation, transformation defaults, the fourteen-test scenario summary, source files and l for "CyFj11: FlowJo v11 Workspace Import and Legacy Format Export for R-Based Flow Cytometry Analysis"

Thirteen synthetic test scenarios were constructed to systematically exercise all supported gate types (rectangle, polygon, ellipsoid, range, Boolean) and transformation types (linear, biexponential, log, arcsinh) in isolation and combination, including hierarchical and Boolean configurations. Each scenario was generated programmatically with the generate_all_tests.R script in flowjo_export_tests/, exported to a FlowJo v10 workspace with export_flowjo10_workspace(), and then opened in FlowJo v10 and FlowJo v11 for count comparison. A fourteenth real-world test used the public FlowSOM example dataset. The development notebook flowjo_export_tests/roundtrip_experiment.Rmd contains the original exploration and detailed visual checks that informed the generator. All test-generation scripts, synthetic data, and exported workspaces described in this appendix are archived in a companion Zenodo repository (<https://doi.org/10.5281/zenodo.20933199>) and are not distributed with the CyFj11 R package itself.

Isolation tests (tests 01–08) are necessary but not sufficient for validating a FlowJo v10 exporter. The FlowJo workspace format interleaves sample definitions, transformation dictionaries, compensation matrices, group nodes, and a recursively nested population tree. Mistakes in how these structures interact are invisible in single-gate tests: transformation parameters must be referenced consistently by channel name at both the <Transformations> and <Axis> levels; gate coordinates must be written in the correct data-space units for the channel’s transform; parent population counts constrain child counts; and Boolean gates are emitted as AndNode, OrNode, or NotNode logical elements whose dependents must resolve to population paths that exist elsewhere in the hierarchy. Tests 09–13 therefore exercise the exporter’s ability to maintain cross-references and coordinate conventions across the whole workspace graph. The small residual count differences observed in these combined tests are consistent with rounding and precision effects in the different mathematical representations used by flowCore and FlowJo, not with systematic exporter errors.

### A.1 Synthetic FCS data generation

For tests 01–13, synthetic FCS files were generated with the helper create_test_fcs() in flowjo_export_tests/generate_all_tests.R, using fixed random seeds to ensure reproducibility. Scatter channels (FSC-A, FSC-H, SSC-A, SSC-H) were drawn from normal distributions (FSC-A: μ = 120,000, σ = 30,000; SSC-A: μ = 80,000, σ = 20,000), and fluorescence channels were drawn from mixtures of two log-normal components (negative population: ~70% of events; positive population: ~30% of events) to produce realistic positive/negative separations. Gate coordinates were chosen to enclose known fractions of the simulated distributions.

### A.2 Transformation defaults

Standard FlowJo-equivalent parameters were used:

| Transformation | Parameters |
| --- | --- |
| Biexponential | channelRange = 4096, maxValue = 262,144, pos = 4.5, neg = 0, widthBasis = -10 |
| Log | decade = 6, offset = 1, scale = 1 |
| Arcsinh (GML2) | T = 262,144, M = 4.5, A = 0 |
| Linear | default minRange = 0, maxRange inferred from FCS $PnR |

### A.3 Test scenario summary

| Test | Purpose | Gate types | Transformations | Hierarchy depth | Notes |
| --- | --- | --- | --- | --- | --- |
| 01 | Root-only baseline | — | Linear | 0 | Verifies minimal workspace export. |
| 02 | Single 1D rectangle gate | Range (1D) on FSC-A [60k, 180k] | Linear | 1 | Tests basic RectangleGate export. |
| 03 | Single 2D rectangle gate | Rect (2D) on FSC-A vs SSC-A | Linear | 1 | Two-dimensional rectangle. |
| 04 | Polygon gate | Polygon (6 vertices) on FSC-A vs FSC-H | Linear | 1 | Singlet-discrimination shape. |
| 05 | Ellipsoid gate | Ellipse on FSC-A vs SSC-A | Linear | 1 | Eigenvalue decomposition on export introduces minor numeric drift. |
| 06 | Biexponential transform | Range (1D) on FITC-A [1000, 3000] | Biexponential | 1 | Tests biex transform serialization. |
| 07 | Log transform | Range (1D) on PE-A [0.4, 0.8] | Log (decade 6) | 1 | Log display-space vs data-space convention; counts still match. |
| 08 | Arcsinh transform | Range (1D) on APC-A [0.25, 0.92] | Arcsinh | 1 | Tests fasinh / GML2 arcsinh serialization. |
| 09 | Two-level hierarchy | Two 2D rectangles | Linear | 2 | cells → singlets. |
| 10 | Boolean AND/OR/NOT | Two 1D ranges + Boolean gates | Linear | 2 | Tests BooleanGate export/import. |
| 11 | Mixed transforms + Boolean | 2D rect, 1D ranges, polygon, Boolean AND/OR | Biexponential (FITC-A), arcsinh (APC-A), linear | 2 | Mixed fluorescence transforms. |
| 12 | Full Boolean set + ellipse | 2D rect, ellipse, ranges, Boolean AND/OR/NOT | Biexponential (FITC-A), log (PE-A), linear | 2 | Includes NOT gate and log drift case. |
| 13 | Deep arcsinh hierarchy | Polygon, 2D rect, ellipse, ranges, Boolean AND/NOT | Arcsinh (FITC-A, PE-A, APC-A), linear | 3 | live → {singlets, FITC_PE_gate} → {FITC_pos, APC_pos, double_pos, not_double}. |
| 14 | Real-world FlowSOM data | Polygon, rectangles | Linear (scatter), biexponential (fluorescence) | 3 | Public mouse bone-marrow dataset (68983.fcs); tests compensation matrix preservation. Populations downstream of the NK1_1+ marker are unavailable in FlowJo v11 because it rewrites stain names containing a slash to an underscore (the FCS $PnS name NK1/1 becomes NK1_1), which breaks the match to the exported gate names NK1_1+ and NK1_1-. |

### A.4 Source files

The authoritative test generator is:

- flowjo_export_tests/generate_all_tests.R — regenerates all 14 test scenarios and writes one FlowJo v10 workspace per scenario (testNN_export.wsp).

A development notebook with detailed visual checks and early parameter exploration is also available:

- flowjo_export_tests/roundtrip_experiment.Rmd — originally used to prototype the round-trip pipeline; not required for reproducing the validation results.

### A.5 File locations

Generated artifacts for each test are stored under flowjo_export_tests/testNN/:

| File | Description |
| --- | --- |
| sampleNN.fcs | Synthetic or real-world FCS file used as input. |
| testNN_export.wsp | FlowJo v10 workspace produced by export_flowjo10_workspace(). |
| testNN_export.llm.wsp | Copy of the exported workspace used for manual comparison and saved from FlowJo v10. |

Test-14 additionally contains the public FlowSOM files 68983.fcs and gating.wsp, plus the CyFj11-exported workspace test14_export.wsp.

### A.6 Validation methodology and acceptance criteria

The validation pipeline and per-population results are described in Appendix S2. The metrics and acceptance criteria used to judge success are summarized below.

| Metric | Acceptance criterion |
| --- | --- |
| Population count difference | ≤ 0.5% absolute |
| Gate coordinate deviation | ≤ 0.1% of axis range (95th percentile) |
| KS D-statistic (transformations) | < 0.05 for ≥ 95% of comparisons |
| Boolean membership agreement | 100% |

### A.7 Regeneration

Tests 01–14 can be regenerated from the root of the Zenodo repository by running:

source("flowjo_export_tests/generate_all_tests.R")

or from the shell:

Rscript flowjo_export_tests/generate_all_tests.R

The script automatically copies 68983.fcs and gating.wsp from the FlowSOM Bioconductor package’s extdata/ directory when FlowSOM is installed; otherwise Test-14 stops with an informative error.

Software environment: All results reported here were produced with CyFj11 v0.99.95 on R 4.5.0 and Bioconductor 3.22 (flowCore 2.22.1, flowWorkspace 4.22.1, openCyto 2.22.0, CytoML 2.24.0, and FlowSOM 2.18.0) under macOS 26.5.2 on Apple Silicon (M3/M4). Small residual count and coordinate differences (Appendix S2) may vary with platform and CPU architecture owing to floating-point rounding.
