## Supplementary material for "CyFj11: FlowJo v11 Workspace Import and Legacy Format Export for R-Based Flow Cytometry Analysis": Per-population validation results, including Table S1 (per-population counts for FlowJo v10 export and FlowJo v11 import across all test scenarios)

### Appendix S2: FlowJo v11 Compatibility and Per-Population Validation

The exported FlowJo v10 workspaces were opened in FlowJo v10 and FlowJo v11, and population counts were recorded at each stage. This appendix documents the observed compatibility behavior of FlowJo v11’s v10 importer and provides the full per-population validation data.

#### B.1 Validation pipeline

For each test scenario:

1. export_flowjo10_workspace() wrote a v10 XML workspace.
2. The workspace was opened in FlowJo v10 and population counts (N_FJ10) were recorded.
3. The workspace was also saved from within FlowJo v10 under “.llm.wsp”.
4. The same workspace was opened in FlowJo v11; counts (N_FJ11) and any abnormalities or discrepancies from the v10 reference were recorded.
5. Where gates were dropped or mis-rendered in FlowJo v11, the gate was manually reconstructed in the v11 GUI and counts were re-recorded (N_FJ11r).
6. The v11-saved workspace was re-read into R via fj11_to_gatingset() to obtain N_re.

Residual differences (res_FJ10, res_FJ11, res_FJ11r) remove the error inherited from the parent population, so they reflect only gate-level discrepancy.

#### B.2 FlowJo v10 re-save effects

Each exported workspace (testNN_export.wsp) was opened and saved in FlowJo v10, producing a corresponding .llm.wsp file. Direct XML comparison shows that FlowJo re-serializes the workspace rather than preserving it verbatim. The most conspicuous changes are cosmetic: clientTimestamp, modDate, and nonAutoSaveFileName are updated; gate IDs are regenerated; lineWeight attributes change from numeric values to enumerated labels such as Hairline; percentX/percentY attributes are dropped; empty <Subpopulations> nodes are removed; and <Graph> subtrees are added under population nodes for group-level plotting. Gate coordinates are also reformatted (e.g., rectangle min/max values acquire a trailing .0, polygon vertices are rounded to integers or fewer decimal places), but the underlying geometry is unchanged.

FlowJo also rewrites the transformation definitions. The CyFj11 export either omits a global <Transformations> block or embeds parameter-level transforms; after a FlowJo save a normalized <Transformations> block appears. Observed rewrites include biex length="4096" → length="256", fasinh M="4" length="1" → M="5.418..." length="256", and log decades="6" → a data-dependent value near 4.1–4.3. These transformations affect how fluorescence channels are displayed, but because FlowJo recomputes populations on load, they do not necessarily change the event counts reported in the GUI.

A handful of stored count attributes in the XML differ between .wsp and .llm.wsp by a few cells (e.g., test 04 singlets: 7,555 → 7,569; test 13 FITC_PE_gate: 2,983 → 2,992). These are below 0.2% and are not user-visible discrepancies: they reflect when FlowJo last recomputed and wrote the count attribute, not a real difference in gated populations. The counts reported in Table S1 were obtained from the FlowJo GUI after loading each workspace, and they are the authoritative values used for validation.

#### B.3 FlowJo v11 compatibility findings

##### B.3.1 Boolean gates

FlowJo v11’s v10 importer **dropped hierarchical Boolean gates** in tests 11–13. Root-level Boolean gates (test 10) were imported, but deeper Boolean populations appeared as “not imported” (N/A) and had to be reconstructed manually. After reconstruction, counts matched the R reference exactly. This behavior localizes the failure to FlowJo v11’s importer, not to the exported XML: the same files open correctly in FlowJo v10.

##### B.3.2 Arcsinh ellipsoid gates

One-dimensional arcsinh rectangle gates matched faithfully between R, FlowJo v10, and FlowJo v11. The **arcsinh ellipsoid gate** in test 13 (FITC_PE_gate on FITC-A vs PE-A) was rendered incorrectly by FlowJo v11 (recorded as 0 events because the gate was effectively dropped), versus the R reference of 2,983 events. FlowJo v10 reproduced the count within numerical noise. This indicates that FlowJo v11’s importer mis-handles the arcsinh-scaled EllipsoidGate coordinates, whereas the v10 XML itself is valid. A likely contributing factor is that FlowJo v11 does not expose a parameterised arcsinh scale at all: its axis-transform panel offers only Linear, Log, Biex, and “ArcTanh”, and the latter is configurable solely through Min and Max bounds, with no user-accessible M, T, W, or length parameters. Consequently the fasinh parameters carried in the v10 workspace (see B.2) cannot be represented faithfully in v11, and gates defined in arcsinh space are re-interpreted against a scale whose parameters v11 has chosen itself.

##### B.3.3 Polygon and ellipse gates (general)

Small residual count differences for polygon gates (test 04, test 11 singlets) and ellipse gates (test 05, test 12 scatter_ellipse) reflect differences in how FlowJo renders these shapes when reading v10-format coordinates, not errors in the gate definitions. Ellipsoid gates are encoded in Gating-ML by two foci and four edge vertices. After re-saving in FlowJo v10 the foci retain full precision but the edge vertices are rounded to integers (e.g., test 05 ellipse_cells: 52.758876... → 52, 48.591041... → 49). This small geometric discretization explains the ≤0.2% drift observed for ellipse gates. Once parent-inherited drift is removed, residuals remain within a few tens of cells (<0.2% relative).

##### B.3.4 Log-transform display space

Test 07 uses a log-transformed PE-A axis. The gate coordinates stored in the v10 XML are in **transformed data space** (0.4–0.8), while FlowJo’s GUI displays them multiplied by the decade parameter (≈6). The counts are nevertheless identical in R and FlowJo, confirming that the data-space coordinates are correct.

##### B.3.5 Real-world test 14 marker-name issue

The test 14 FCS file stores the PE-A stain as NK1/1 and the AmCyan-A stain as l/d (Live/Dead) in the $PnS keywords. FlowJo v11 rewrites stain names that contain a slash (/), replacing it with an underscore, so NK1/1 becomes NK1_1. The exported gate/population names NK1_1+ and NK1_1- then no longer match the rewritten marker name, and all populations gated downstream of NK1_1+ and NK1_1- therefore displayed as not available in FlowJo v11. FlowJo v10 kept the slash and reproduced the NK1_1+ count correctly (residual = 1 event). This failure is a FlowJo v11 stain-name normalization idiosyncrasy and does not indicate an error in the CyFj11 export.

#### B.4 Table S1. Per-population event-count validation

Table S1 gives the full per-population counts for all 14 tests. Columns are defined as follows:

| Column | Meaning |
| --- | --- |
| N_R | Reference count from R (flowCore/openCyto). |
| N_FJ10 | Count in FlowJo v10 after workspace import. |
| N_FJ11 | Count in FlowJo v11 after workspace import. |
| N_FJ11r | Count after gate remade in FlowJo v11. |
| N_re | Count from re-reading the FlowJo v11-exported workspace. |
| res_FJ10 | Residual N_R − N_FJ10 after removing parent-inherited error. |
| res_FJ11 | Residual N_R − N_FJ11 after removing parent-inherited error. |
| res_FJ11r | Residual N_re − N_FJ11r after removing parent-inherited error. |

Abbreviations: **Range (1D)**, one-dimensional range gate; **Rect (2D)**, two-dimensional rectangle gate; **Poly**, polygon gate; **Ellipse**, ellipsoid gate; **AND / OR / NOT**, Boolean gates; **NI**, not imported; **Nim**, not implemented; **N/A**, not evaluated; **—**, not computable.

##### B.4.1 Tests 01–09: single gate-type validation

| Test | Gate | Gate type | Channel(s) | N_R | N_FJ10 | N_FJ11 | N_FJ11r | N_re | res_FJ10 | res_FJ11 | res_FJ11r |
| --- | --- | --- | --- | --- | --- | --- | --- | --- | --- | --- | --- |
| 01 | All Events | Root | — | 5000 | 5000 | 5000 | 5000 | 5000 | 0 | 0 | 0 |
| 02 | FSC_filter | Range (1D) | FSC-A | 4776 | 4776 | 4776 | 4776 | 4776 | 0 | 0 | 0 |
| 03 | cells | Rect (2D) | FSC-A vs SSC-A | 4631 | 4631 | 4631 | 4631 | 4631 | 0 | 0 | 0 |
| 04 | singlets | Poly | FSC-A vs FSC-H | 7555 | 7569 | 7506 | 7506 | 7555 | -14 | 49 | 49 |
| 05 | ellipse_cells | Ellipse | FSC-A vs SSC-A | 7301 | 7306 | 0 | 7290 | 7281 | -5 | 7301 | -9 |
| 06 | FITC_pos | Range (1D) | FITC-A (Biexp) | 2403 | 2403 | 2403 | 2400 | 2400 | 0 | 0 | 0 |
| 07 | PE_pos | Range (1D) | PE-A (log) | 3198 | 3198 | 3198 | 3198 | 3198 | 0 | 0 | 0 |
| 08 | APC_pos | Range (1D) | APC-A (Arcsinh) | 1606 | 1606 | 1608 | 1599 | 1599 | 0 | -2 | 0 |
| 09 | cells | Rect (2D) | FSC-A vs SSC-A | 9441 | 9441 | 9441 | 9441 | 9441 | 0 | 0 | 0 |
| 09 | singlets | Rect (2D) | FSC-A vs FSC-H | 8800 | 8800 | 8800 | 8800 | 8800 | 0 | 0 | 0 |

† Test 05: round-trip count drift is expected for ellipsoid gates because different mathematical representations of the ellipse accumulate rounding and precision effects. In this test, FlowJo v11 could not read the ellipse gate even from the file re-saved by FlowJo v10: it misinterpreted the ellipsoid boundary and crashed, so N_FJ11 is recorded as 0.

##### B.4.2 Test 10: Boolean AND / OR / NOT (linear axes)

| Gate | Path | Gate type | Condition | N_R | N_FJ10 | N_FJ11 | N_FJ11r | N_re | res_FJ10 | res_FJ11 | res_FJ11r |
| --- | --- | --- | --- | --- | --- | --- | --- | --- | --- | --- | --- |
| FSC_gate | /FSC_gate | Range (1D) | FSC-A | 7554 | 7554 | 7554 | 7558 | 7558 | 0 | 0 | 0 |
| SSC_gate | /SSC_gate | Range (1D) | SSC-A | 7392 | 7392 | 7392 | 7392 | 7392 | 0 | 0 | 0 |
| both | /both | AND | FSC_gate and SSC_gate | 6979 | 6979 | 6979 | 6983 | 6983 | 0 | 0 | 0 |
| bothOR | /bothOR | OR | FSC_gate or SSC_gate | 7967 | 7967 | 7967 | 7967 | 7967 | 0 | 0 | 0 |
| bothNOT | /bothNOT | NOT | Not both | 1021 | 1021 | 1021 | 1017 | 1017 | 0 | 0 | 0 |

##### B.4.3 Test 11: mixed biexponential + arcsinh, hierarchy + Boolean

| Gate | Path | Gate type | Channel(s) | N_R | N_FJ10 | N_FJ11 | N_FJ11r | N_re | res_FJ10 | res_FJ11 | res_FJ11r |
| --- | --- | --- | --- | --- | --- | --- | --- | --- | --- | --- | --- |
| cells | /cells | Rect (2D) | FSC-A vs SSC-A | 9696 | 9696 | 9696 | 9696 | 9696 | 0 | 0 | 0 |
| FITC_pos | /cells/FITC_pos | Range (1D) | FITC-A (Biexp) | 2897 | 2897 | 2897 | 2897 | 2897 | 0 | 0 | 0 |
| APC_pos | /cells/APC_pos | Range (1D) | APC-A (Arcsinh) | 1930 | 1930 | 1930 | 1930 | 1930 | 0 | 0 | 0 |
| singlets | /cells/singlets | Poly | FSC-A vs FSC-H | 9478 | 9474 | 9487 | 9478 | 9478 | 4 | -9 | 0 |
| double_pos | /cells/double_pos | AND | FITC_pos and APC_pos | 1928 | 1928 | N/A | 1928 | 1928 | 0 | — | 0 |
| either_pos | /cells/either_pos | OR | FITC_pos or APC_pos | 2899 | 2899 | N/A | 2899 | 2899 | 0 | — | 0 |

##### B.4.4 Test 12: biexponential + log, ellipse + full Boolean set

**Note.** FlowJo v11 does not support a combined NOT(pop1, pop2) Boolean gate. PE_hi and gates depending on it may exhibit round-trip count drift attributable to the log transform. In this test FlowJo v11 also failed to read the scatter_ellipse ellipsoid gate from the re-saved v10 file, misinterpreted the boundary, and crashed, so N_FJ11 is recorded as 0.

| Gate | Path | Gate type | Channel(s) | N_R | N_FJ10 | N_FJ11 | N_FJ11r | N_re | res_FJ10 | res_FJ11 | res_FJ11r |
| --- | --- | --- | --- | --- | --- | --- | --- | --- | --- | --- | --- |
| cells | /cells | Rect (2D) | FSC-A vs SSC-A | 9765 | 9765 | 9765 | 9765 | 9765 | 0 | 0 | 0 |
| scatter_ellipse | /scatter_ellipse | Ellipse | FSC-A vs SSC-A | 9803 | 9797 | 0* | 9768 | 9772 | 6 | 9803 | 4 |
| FITC_hi | /cells/FITC_hi | Range (1D) | FITC-A (Biexp) | 2934 | 2934 | 2934 | 2934 | 2934 | 0 | 0 | 0 |
| PE_hi | /cells/PE_hi | Range (1D) | PE-A (log) | 3831 | 3831 | 3831 | 3831 | 3831 | 0 | 0 | 0 |

* N_FJ11 = 0 because FlowJo v11 crashed when reading the scatter_ellipse ellipsoid gate from the v10-re-saved file. | double_hi | /cells/double_hi | AND | FITC_hi and PE_hi | 2879 | 2879 | N/A | 2879 | 2879 | 0 | — | 0 | | either_hi | /cells/either_hi | OR | FITC_hi or PE_hi | 3886 | 3886 | N/A | 3886 | 3886 | 0 | — | 0 | | neither | /cells/neither | NOT | Not either_hi | 5879 | 5879 | N/A | 5934 | N/A | 0 | — | — |

##### B.4.5 Test 13: arcsinh × 3 channels, 3-level hierarchy + Boolean

**Note.** As in tests 05 and 12, FlowJo v11 failed to read the arcsinh ellipsoid gate (FITC_PE_gate) from the v10-re-saved file and crashed, so N_FJ11 is recorded as 0.

| Gate | Path | Gate type | Channel(s) | N_R | N_FJ10 | N_FJ11 | N_FJ11r | N_re | res_FJ10 | res_FJ11 | res_FJ11r |
| --- | --- | --- | --- | --- | --- | --- | --- | --- | --- | --- | --- |
| live | /live | Poly | FSC-A vs SSC-A | 9939 | 9940 | 9938 | N/A | N/A | -1 | 1 | — |
| singlets | /live/singlets | Rect (2D) | FSC-A vs FSC-H | 9470 | 9470 | 9470 | N/A | N/A | 1 | -1 | — |
| FITC_PE_gate | /live/FITC_PE_gate | Ellipse | FITC-A vs PE-A (Arcsinh) | 2983 | 2992 | 0* | N/A | N/A | -9 | 2983 | — |
| FITC_pos | /live/singlets/FITC_pos | Range (1D) | FITC-A (Arcsinh) | 2838 | 2838 | 2838 | N/A | N/A | 0 | 0 | — |
| APC_pos | /live/singlets/APC_pos | Range (1D) | APC-A (Arcsinh) | 1888 | 1888 | 1888 | N/A | N/A | 0 | 0 | — |

* N_FJ11 = 0 because FlowJo v11 crashed when reading the arcsinh ellipsoid gate from the v10-re-saved file. | double_pos | /live/singlets/double_pos | AND | FITC_pos and APC_pos | 1887 | 1887 | N/A | N/A | N/A | 0 | — | — | | not_double | /live/singlets/not_double | NOT | Not double_pos | 7583 | 7583 | N/A | N/A | N/A | 0 | — | — |

##### B.4.6 Test 14: real-world validation — FlowSOM example data

**Source.** FlowSOM R package; file 68983.fcs + gating.wsp (mouse bone marrow immunophenotyping).
**Pipeline.** CytoML::flowjo_to_gatingset() → export_flowjo10_workspace() (CyFj11 v10 → CytoML v10 import).
**Acquisition settings.** Compensation matrix applied; fluorescence channels biexponential; scatter channels linear. **Round-trip check.** N_FJ10₂ / N_FJ11₂: counts after the FlowJo v11–corrected workspace (the source of N_re) was re-exported a second time via export_flowjo10_workspace() and reopened in FlowJo v10 and v11, testing round-trip stability of the export function and whether the NK1_1 marker-name issue affecting N_FJ11 persists after correction. All 15 populations, including those previously not available (N/A) in N_FJ11, were read without error in this second round.

| Gate | Gate type | Channel(s) | N_R | N_FJ10 | N_FJ11 | N_FJ11r | N_re | res_FJ10 | res_FJ11 | res_FJ11r | N_FJ10₂ | N_FJ11₂ |
| --- | --- | --- | --- | --- | --- | --- | --- | --- | --- | --- | --- | --- |
| Lymphocytes | Poly | FCS-SSC | 19212 | 19203 | 19132 | 19132 | 19079 | 9 | 80 | -53 | 19071 | 19132 |
| Singlets | Rect (2D) | FSC-H FSC-W | 18689 | 18680 | 18594 | 18594 | 18545 | 9 | 95 | -49 | 18537 | 18594 |
| Singlets2 | Rect (2D) | SSC-H SSC-W | 18573 | 18563 | 18466 | 18466 | 18418 | 10 | 107 | -48 | 18410 | 18466 |
| Nk1_1+ | Rect (2D) | NK1.1 vs FSC-A | 923 | 922 | N/A | 920 | 942 | 1 | — | 22 | 942 | 939 |
| NK cells | Rect (2D) | CD3 NK1.1 | 312 | 312 | N/A | 312 | 314 | 0 | — | 2 | 314 | 315 |
| NK T cells | Rect (2D) | CD3 NK1.1 | 535 | 534 | N/A | 533 | 545 | 1 | — | 12 | 546 | 543 |
| NK1_1- | Rect (2D) | NK1.1 vs FSC-A | 17458 | 17449 | N/A | 17357 | 17192 | 9 | — | -165 | 17184 | 17240 |
| B cells | Rect (2D) | CD19 CD3 | 2459 | 2460 | N/A | 2452 | 2436 | -1 | — | -16 | 2434 | 2431 |
| T cells | Rect (2D) | CD19 CD3 | 10745 | 10747 | N/A | 10716 | 10672 | -2 | — | -44 | 10669 | 10652 |
| Ab T cells | Poly | TCRβ vs TCRγδ | 9188 | 9190 | N/A | 9165 | 9113 | -2 | — | -52 | 9112 | 9096 |
| CD4 T cells | Rect (2D) | CD4 vs CD8 | 7480 | 7485 | N/A | 7462 | 7411 | -5 | — | -51 | 7410 | 7394 |
| CD8 T cells | Rect (2D) | CD4 vs CD8 | 1407 | 1404 | N/A | 1407 | 1403 | 3 | — | -4 | 1403 | 1403 |
| DN T cells | Rect (2D) | CD4 vs CD8 | 266 | 266 | N/A | 261 | 263 | 0 | — | 2 | 263 | 261 |
| DP T cells | Rect (2D) | CD4 vs CD8 | 6 | 6 | N/A | 6 | 7 | 0 | — | 1 | 7 | 7 |
| Gd T cells | Poly | TCRβ vs TCRγδ | 1470 | 1470 | N/A | 1466 | 1463 | 0 | — | -3 | 1464 | 1464 |

**[a]** The original FlowSOM workspace was written by FlowJo 10.5.3. FlowJo 11.2 was unable to read this file and did not correctly interpret the biexponential-transformed channels. Re-saving the workspace in FlowJo 10.10.1 produced a file that FlowJo 11.2 could open.

All other discrepancies can be explained by numerical conversion differences.

##### B.4.7 Test 14 round-trip re-verification (second export cycle)

| Gate | N_FJ11r | N_re | N_FJ10₂ | N_FJ11₂ |
| --- | --- | --- | --- | --- |
| Lymphocytes | 19132 | 19079 | 19071 | 19132 |
| Singlets | 18594 | 18545 | 18537 | 18594 |
| Singlets2 | 18466 | 18418 | 18410 | 18466 |
| Nk1_1+ | 920 | 942 | 942 | 939 |
| NK cells | 312 | 314 | 314 | 315 |
| NK T cells | 533 | 545 | 546 | 543 |
| NK1_1- | 17357 | 17192 | 17184 | 17240 |
| B cells | 2452 | 2436 | 2434 | 2431 |
| T cells | 10716 | 10672 | 10669 | 10652 |
| Ab T cells | 9165 | 9113 | 9112 | 9096 |
| CD4 T cells | 7462 | 7411 | 7410 | 7394 |
| CD8 T cells | 1407 | 1403 | 1403 | 1403 |
| DN T cells | 261 | 263 | 263 | 261 |
| DP T cells | 6 | 7 | 7 | 7 |
| Gd T cells | 1466 | 1463 | 1464 | 1464 |

#### B.5 Test 14 detail: compensation matrix preservation

Test 14 exercises compensation matrix round-tripping on real mouse bone-marrow data. The original FlowSOM workspace (flowjo_export_tests/test14/test14_flowsom.wsp) contains an 11 × 11 acquisition-defined spillover matrix. CyFj11 exports this matrix in flowjo_export_tests/test14/test14.compensation.wsp. The original matrix stores coefficients as proportions (diagonal = 1), while the exported matrix stores them as percentages (diagonal = 100); after rescaling the exported matrix by ÷100, the element-wise comparison is:

- Matrix dimension: 11 × 11
- Mean absolute relative difference: 0.000308%
- Median absolute relative difference: 0.000010%
- Maximum absolute relative difference: 0.011125% (Qdot 605-A → APC-Cy7-A)

The full element-wise comparison is provided in manuscript/test14_compensation_elementwise.csv; the 20 largest absolute relative differences are shown in Table B2.

##### Table B2. Largest absolute relative differences in the test 14 compensation matrix

| From | To | Original | Exported (scaled) | Absolute diff | Relative diff (%) |
| --- | --- | --- | --- | --- | --- |
| Qdot 605-A | APC-Cy7-A | 0.0000261 | 0.0000261 | 2.9e-09 | 0.011125 |
| Qdot 605-A | PE-Cy7-A | 0.0000364 | 0.0000364 | 2.7e-09 | 0.007414 |
| Pacific Blue-A | Alexa Fluor 700-A | 0.0001085 | 0.0001085 | -2.1e-09 | -0.001935 |
| Pacific Blue-A | PE-Cy7-A | 0.0000388 | 0.0000387 | -7.0e-10 | -0.001806 |
| Qdot 605-A | Pacific Blue-A | 0.0002496 | 0.0002496 | -4.4e-09 | -0.001763 |
| Qdot 605-A | Alexa Fluor 700-A | 0.0002576 | 0.0002576 | -4.5e-09 | -0.001747 |
| Qdot 605-A | FITC-A | 0.0000863 | 0.0000863 | -1.5e-09 | -0.001739 |
| AmCyan-A | PE-Texas Red-A | 0.0001757 | 0.0001757 | 2.6e-09 | 0.001480 |
| Pacific Blue-A | FITC-A | 0.0004844 | 0.0004844 | -4.4e-09 | -0.000908 |
| PE-Texas Red-A | Alexa Fluor 700-A | 0.0001830 | 0.0001830 | 1.1e-09 | 0.000601 |
| PE-Texas Red-A | Pacific Blue-A | 0.0005403 | 0.0005403 | -2.7e-09 | -0.000500 |
| FITC-A | APC-Cy7-A | 0.0008660 | 0.0008660 | 3.8e-09 | 0.000439 |
| PE-A | FITC-A | 0.0011664 | 0.0011664 | -4.9e-09 | -0.000420 |
| PE-Cy5-A | Pacific Blue-A | 0.0005018 | 0.0005018 | -1.7e-09 | -0.000339 |
| AmCyan-A | Alexa Fluor 700-A | 0.0011353 | 0.0011353 | 3.8e-09 | 0.000335 |
| PE-A | Alexa Fluor 700-A | 0.0002449 | 0.0002449 | -8.0e-10 | -0.000327 |
| PE-Cy5-A | FITC-A | 0.0003372 | 0.0003372 | 1.1e-09 | 0.000326 |
| PE-Cy5-A | AmCyan-A | 0.0016727 | 0.0016727 | 4.9e-09 | 0.000293 |
| FITC-A | Alexa Fluor 700-A | 0.0010697 | 0.0010697 | -3.0e-09 | -0.000280 |
| Pacific Blue-A | APC-A | 0.0017360 | 0.0017360 | -3.8e-09 | -0.000219 |

The compensation matrix is therefore preserved to numerical round-off; remaining differences are at the level of single-precision representation and do not affect population counts.

#### B.6 Summary statistics

Key summary statistics derived from Table S1 are reproduced in Table B3.

##### Table B3. Validation summary by comparison subset

| Comparison subset | N | Exact match | Within ≤0.3% | Max | residual |
| --- | --- | --- | --- | --- | --- |
| Export FJ10 | synthetic, all gates | 35 | 28 (80%) | 7 (20%) | 14 cells |
| Export FJ10 | synthetic core (excl. arcsinh+bool) ★ | 20 | 14 (70%) | 6 (30%) | 14 cells |
| Export FJ10 | arcsinh gates only | 5 | 4 (80%) | 1 (20%) | 9 cells |
| Export FJ10 | boolean gates only | 10 | 10 (100%) | 0 (0%) | 0 cells |
| Export FJ10 | real-world test14 | 15 | 4 (27%) | 11 (73%) | 10 cells |
| Import R | synthetic, all gates | 27 | 24 (89%) | 1 (4%) | 73 cells |
| Import R | synthetic core (excl. arcsinh+bool) ★ | 18 | 15 (83%) | 1 (6%) | 73 cells |
| Import R | arcsinh gates only | 2 | 2 (100%) | 0 (0%) | 0 cells |
| Import R | boolean gates only | 7 | 7 (100%) | 0 (0%) | 0 cells |
| Import R | real-world test14 | 15 | 1 (7%) | 7 (47%) | 90 cells |

The synthetic export and import comparisons pass the acceptance criterion of mean absolute percentage difference ≤0.5%. The real-world test 14 import comparison exceeds this threshold because FlowJo v11 rewrites the NK1/1 stain name to NK1_1, breaking the gate names NK1_1+ and NK1_1-, and because of small residual drift accumulated across a deep compensation/transform stack.

#### B.7 Gating-ML 2.0 conformance

Structural Gating-ML 2.0 conformance of the exported v10 workspaces is documented in Appendix S3. All 50 applicable structural checks passed across the 13 synthetic test workspaces (100%). That appendix also contains the per-test conformance table (Table C1).

#### B.8 Technical implementation notes

Details of the FlowJo v10/v11 format differences, gate-type conversions, transformation parameter mapping, package architecture, and validation methodology are provided in Appendix S4.
