## Supplementary material for "CyFj11: FlowJo v11 Workspace Import and Legacy Format Export for R-Based Flow Cytometry Analysis": Gating-ML 2.0 standards-compliance results: the 50 applicable structural checks across the 13 test workspaces

### Appendix S3: Gating-ML 2.0 Standards Compliance

CyFj11 exports FlowJo v10 workspaces as XML documents intended to conform to the ISAC Gating-ML 2.0 standard (Spidlen et al., *Cytometry A*, 2012). This appendix documents the structural conformance checks performed on the 13 synthetic test workspaces and summarizes the results.

#### S3.1 Conformance check methodology

The R script manuscript/validate_gatingml_conformance.R performs structural checks on each exported workspace without requiring network access to the published XSD; the structural rules are implemented directly in the script so that the check is self-contained and reproducible offline. The checks mirror the structural rules defined in the published Gating-ML 2.0 specification and XSD:

1. **Namespace declarations.** The Gating-ML 2.0 namespace URIs for gating, transformations, and data-type must be declared.
2. **Gate element structure.** Each RectangleGate, PolygonGate, and EllipsoidGate must contain the required child elements and attributes, and every dimension must reference a named FCS parameter.
3. **Transformation element structure.** Each linear, biex, fasinh (arcsinh), and log transformation element must carry the required parameter attributes and a non-empty data-type:parameter child.

Checks are run only on SampleNode elements to avoid duplicate counting of group-level gate definitions.

#### S3.2 Results

All 50 applicable structural checks passed across the 13 synthetic test workspaces (100%). The per-test breakdown is shown in Table S3.1.

#### S3.3 Table S3.1. Gating-ML 2.0 structural conformance by test

| Test | Namespace | Linear transform | RectangleGate | PolygonGate | EllipsoidGate | Biex transform | Log transform | Arcsinh transform |
| --- | --- | --- | --- | --- | --- | --- | --- | --- |
| test01 | ✓ | ✓ | — | — | — | — | — | — |
| test02 | ✓ | ✓ | ✓ | — | — | — | — | — |
| test03 | ✓ | ✓ | ✓ | — | — | — | — | — |
| test04 | ✓ | ✓ | — | ✓ | — | — | — | — |
| test05 | ✓ | ✓ | — | — | ✓ | — | — | — |
| test06 | ✓ | ✓ | ✓ | — | — | ✓ | — | — |
| test07 | ✓ | ✓ | ✓ | — | — | — | ✓ | — |
| test08 | ✓ | ✓ | ✓ | — | — | — | — | ✓ |
| test09 | ✓ | ✓ | ✓ | — | — | — | — | — |
| test10 | ✓ | ✓ | ✓ | — | — | — | — | — |
| test11 | ✓ | ✓ | ✓ | ✓ | — | ✓ | — | ✓ |
| test12 | ✓ | ✓ | ✓ | — | ✓ | ✓ | ✓ | — |
| test13 | ✓ | ✓ | ✓ | ✓ | ✓ | — | — | ✓ |
| **Total passing / applicable** | **13 / 13** | **13 / 13** | **10 / 10** | **3 / 3** | **3 / 3** | **3 / 3** | **2 / 2** | **3 / 3** |

Legend: ✓ = check passed; — = gate/transform type not present in that test.

#### S3.4 Coverage notes

The structural checks verify conformance with the structural rules of the Gating-ML 2.0 schema, not semantic correctness of population counts. Because the rules are reimplemented in the check script rather than applied by an XML validator, they are not a substitute for full XSD validation against the published schema. Semantic correctness is assessed separately in Appendix S2. Details of the export implementation — including the FlowJo v10/v11 format comparison, gate-type conversions, and transformation parameter mapping — are provided in Appendix S4. The two checks are complementary:

- **Structural conformance indicates that exported workspaces should be parseable by a** Gating-ML 2.0–compliant reader.
- **Population-count validation** establishes that the gates encoded in those XML structures produce the expected event subsets for the test scenarios examined.

#### S3.5 Reproducibility

The conformance check can be re-run from the package root with:

Rscript manuscript/validate_gatingml_conformance.R

This regenerates manuscript/gatingml_conformance.csv and prints the per-test report shown above.
