## Supplementary material for "CyFj11: FlowJo v11 Workspace Import and Legacy Format Export for R-Based Flow Cytometry Analysis": Technical implementation details: FlowJo v10/v11 workspace format comparison, gate-type conversion, transformation parameter mapping, and package arch

### Appendix S4: Technical Implementation Details

This appendix describes the internal design of CyFj11 for readers interested in how the package maps between the FlowJo v11 JSON workspace, R/Bioconductor GatingSet objects, and the FlowJo v10 XML format. It covers the two workspace formats, gate-type conversions, transformation parameter mapping, and the organization of the R source code.

#### S4.1 FlowJo workspace format comparison

FlowJo v10 and v11 use fundamentally different file formats. The design of CyFj11 is shaped by these differences (Table S4.1).

Table S4.1. Comparison of the FlowJo v10 and FlowJo v11 workspace formats

| Aspect | FlowJo v10 (.wsp) | FlowJo v11 (.flowjo) |
| --- | --- | --- |
| File format | Single XML file | ZIP archive containing multiple JSON files |
| Standard | ISAC Gating-ML 2.0 (published) | Proprietary (undocumented) |
| Gate representation | Nested XML elements under each population | UUID-linked JSON objects in populationDefinitions |
| Transformation storage | Separate transforms:* elements | Embedded in axis parameter specifications |
| Per-sample gate variation | Not supported | desyncTable mechanism for tailored gates |
| Population hierarchy | Nested XML containment | Parent/child UUID references |
| Cross-references | Self-contained populations | Global UUID graph across ~10 entity types (groups, dataSources, populationDefinitions, populations, compoundParameterSets, compoundPopulations, paramsetDefinitions, platforms, cytometers, analysisRoot) |

Because the v11 JSON schema is proprietary and not publicly documented, CyFj11 implements **import of the v11 JSON structures that are most commonly needed in practice** by reverse-engineering them, but targets **v10 XML for export**. The implemented import covers the gate types, transformations, compensation matrices, sample groups, and per-sample gate variations that account for the majority of real-world gating workflows, but it does not claim completeness for every FlowJo v11 feature. A practical consideration reinforced the decision to export to v10 rather than write native .flowjo files: FlowJo v11 gives no informative error messages when it fails to load a workspace, the outcome is not reproducible even when reading the same file twice, and successful opening can depend on clearing the application cache or restarting the BD software security layer. The v10 XML path avoids this opaque environment because FlowJo v11 can open v10-format workspaces directly and because any structural problem in the XML can be diagnosed with standard validation tools. In addition, FlowJo v11 is still under active development; rather than continuously chasing an evolving, undocumented binary format, we expect that a mature FlowJo v11 importer for the well-documented v10 XML standard will ultimately be FlowJo’s own responsibility. This asymmetric design therefore provides a practical, maintainable interoperability path without requiring an unsupported write-side reverse-engineering of the v11 schema.

#### S4.2 Gate type conversion

##### S4.2.1 Rectangle gates

v11 rectangle gates define min/max boundaries per parameter. On import these are converted to flowCore::rectangleGate objects. On export to v10 they become <gating:RectangleGate> elements with <gating:dimension> children carrying gating:min and gating:max attributes.

One-dimensional rectangle gates (called range gates in v11 terminology) are imported as rectangleGate objects with one bounded dimension and the other dimension spanning -Inf to +Inf. Neither FlowJo v10 nor flowCore defines a distinct range-gate type; a range gate is represented as a rectangle gate with only one constrained dimension. On export to v10 CyFj11 therefore writes the same <gating:RectangleGate> structure with a single <gating:dimension>.

##### S4.2.2 Polygon gates

v11 polygon gates store vertex coordinates as arrays of [x, y] pairs. On import, vertices are converted to a flowCore::polygonGate matrix. Coordinate inverse-transformation is applied to map from FlowJo display space to raw data space. On export, vertices are written as <gating:vertex> elements inside <gating:PolygonGate>.

##### S4.2.3 Ellipsoid gates

v11 ellipsoid gates are parameterized by a mean vector, a covariance matrix, and a Mahalanobis distance. flowCore uses the same parameterization (ellipsoidGate objects store mean, cov, and distance), so on import CyFj11 maps the v11 ellipse parameters directly to a flowCore::ellipsoidGate without changing the underlying mathematical representation. On export to v10, the covariance matrix is decomposed to recover the rotation angle and semi-axis lengths required by FlowJo v10’s <gating:EllipsoidGate> encoding (which stores foci and edge points rather than covariance). This eigenvalue decomposition introduces minor numerical differences relative to the original v11 gate, which is expected and observed in the validation results.

##### S4.2.4 Boolean gates

v11 Boolean gates reference other gates by UUID and specify AND, OR, or NOT combinations. In the v11 JSON structure these Boolean populations are stored separately from the main gating hierarchy and must be processed only after all referenced concrete gates have been resolved. This two-phase requirement — first material gates, then logical combinations — makes Boolean handling one of the more fragile parts of the import pipeline. On import, CyFj11 resolves UUID references to population paths and constructs a flowCore::booleanFilter. On export, Boolean logic is preserved in <gating:BooleanGate> elements that reference dependent gates by population path.

##### S4.2.5 Quadrant gates

v11 quadrant gates define dividing lines on two axes, producing four quadrant populations. On import, CyFj11 converts each quadrant into a flowCore::quadGate object. On export to v10, quadrant gates are not currently implemented: CyFj11’s exporter handles rectangle, polygon, ellipsoid, and Boolean gates, and a quadGate object is therefore skipped with a warning. This is a current limitation of the CyFj11 exporter rather than a definitive statement about the v10 XML format, which can represent dividing lines via <gating:QuadrantGate> elements in principle.

#### S4.3 Transformation parameter mapping

All transformations are extracted from v11 axis definitions and converted to the corresponding flowCore transformation objects on import. On export to v10 they are written as <transforms:*> elements with the appropriate parameter attributes.

##### S4.3.1 Display space vs. data space

An important distinction when working with FlowJo transformations is that FlowJo displays gate coordinates in **display space** (the transformed, scaled axis values shown on screen), while flowCore and the Gating-ML 2.0 standard store gate coordinates in **data space** (raw channel values). By default CyFj11 converts FlowJo display-space coordinates back to raw data space during import by applying the inverse transformation. Each inverse operation is a mathematical approximation, so converting many vertices or ellipse parameters back and forth can accumulate small rounding errors.

To give users control over this trade-off, CyFj11 exposes two import parameters:

- **use_transformed_coords** — if TRUE, gate coordinates are kept in transformed (display) space instead of being mapped back to raw data space. This avoids inverse-transform rounding but means the resulting flowCore gate operates on transformed data, which is not the default expectation of downstream tools.
- **correct_faulty_gate** — a fallback numeric value used when a transform reports maxRange = 0, which would otherwise make scaling undefined. It allows the import to proceed for workspaces with malformed linear-range metadata.

On export the forward transformation is applied, so coordinates written to the v10 XML are in data space. Users inspecting gate coordinates directly in the v10 XML should be aware that these are in data space and will differ from the values shown on FlowJo’s axis. Counts are nevertheless reproduced correctly because the data-space values are consistent.

##### S4.3.2 Parameter mapping table

Table S4.2 lists the transformations that FlowJo v11 can store and how they map to the parameters used in flowCore functions and in FlowJo v10 XML attributes. The flowCore names are the arguments of the corresponding transform constructor; the v10 XML names are the attributes written by export_flowjo10_workspace().

Table S4.2. Transformation parameter mapping between FlowJo v11, flowCore and FlowJo v10 XML

| Transformation | v11 parameter names | flowCore constructor parameters | v10 XML attribute names | Notes |
| --- | --- | --- | --- | --- |
| Biexponential | T, W, M, A | biexponentialTransform(): a, b, c, d, f, w | T, W, M, A | FlowJo’s T/W/M/A are computed from the flowCore biexponential coefficients and written as v10 attributes |
| Log | decades, offset | logTransform(): logbase, r, d | decades, offset | Base 10 assumed |
| Arcsinh | T, A (stored in v11 as a, b, c) | arcsinhTransform(): a, b, c | length, maxRange, T, A, M, W | v10 XML stores the GML2-style fasinh parameterization; flowCore uses the simpler asinh(a+b*x)+c form |
| Linear | min, max | linearTransform(): a, b | minRange, maxRange | maxRange inferred from FCS $PnR when absent |

Logicle is supported by flowCore (logicleTransform(): w, t, m, a) and can in principle be written to v10 XML, but FlowJo v11 does not expose Logicle as a user-selectable transformation. It therefore does not appear in the v11 → v10 mapping path covered by the validation suite.

The Arcsinh entry warrants additional comment. FlowJo v11 stores and reads arcsinh parameters internally, but its GUI renders arcsinh axes with an arctanh view and does not expose the underlying arcsinh parameters for editing. Consequently, arcsinh-based gates cannot be reliably verified or adjusted inside FlowJo v11, even when the exported XML itself is mathematically correct. This limitation affects all tools that rely on FlowJo v11’s v10-importer path, not only CyFj11.

#### S4.4 Compensation channel naming

When compensation matrices extracted from a FlowJo v11 workspace were re-exported to FlowJo v10 XML, compensation information could be lost, leaving the exported workspace unreadable by FlowJo. Diagnosing this failure uncovered a previously undocumented quirk in how FlowJo names compensated channels, described below together with CyFj11’s handling of it.

##### S4.4.1 Root cause

FlowJo v11 and the flowCore/flowWorkspace ecosystem apply four distinct, mutually inconsistent conventions to compensated channel names, and CyFj11 must reconcile all four to preserve compensation metadata across the round trip:

**FlowJo v11 storage** — compensation matrices store channel names with a Comp- prefix in the parameters list (e.g., Comp-FITC-A), not the underlying detector name (e.g., FITC-A).

**flowCore/flowWorkspace expectation** — channels produced by flowCore::compensate() must carry a Comp- prefix to be recognised by downstream FlowJo export functions.

**FlowJo export requirement** — the FCS header $PnN keywords for compensated channels must carry a double Comp-Comp- prefix (e.g., Comp-Comp-FITC-A) before FlowJo v10 and v11.2 will read the exported workspace; this is a FlowJo parsing requirement rather than documented behaviour.

**Gate dimension naming** — gates reference compensated channels with a single Comp- prefix (e.g., Comp-FITC-A), consistent with how flowWorkspace stores compensated channel names internally.

None of these conventions is documented by FlowJo, and the double Comp-Comp- requirement for $PnN keywords may reflect a parsing idiosyncrasy in FlowJo v10/v11 that users must work around for interoperability.

##### S4.4.2 CyFj11’s solution

CyFj11 reconciles these four naming layers explicitly rather than assuming a single consistent convention. The package:

Extracts compensation matrices using the Comp-prefixed channel names from FlowJo v11’s parameters list.

Renames channels to Comp-<channel> after applying compensation.

Writes $PnN keywords with a Comp-Comp-<channel> prefix on export, for FlowJo compatibility.

Writes gate dimensions with a single Comp-<channel> prefix.

##### S4.4.3 Impact

This reconciliation is what allows compensation metadata to survive the full FlowJo v11 → v10 → v11 round trip described in S4.1, underpinning the entry-by-entry compensation-matrix fidelity reported for Test 14 in the main text and supporting reproducible cytometry analysis workflows more generally.

#### S4.5 Package architecture

CyFj11 is organized into the following R source modules (Table S4.3).

Table S4.3. CyFj11 R source modules and their responsibilities

| File | Responsibility |
| --- | --- |
| R/archive.R | ZIP extraction, JSON parsing, manifest reading |
| R/conversion.R | Per-sample GatingSet list construction from parsed workspace |
| R/export-flowjo10.R | Gating-ML 2.0 XML generation |
| R/gates.R | Gate type detection, coordinate extraction, conversion |
| R/transformations.R | Transformation extraction and format conversion |
| R/compensation.R | Compensation matrix extraction and validation |
| R/populations.R | Population hierarchy traversal and gate tree construction |
| R/file-search.R | FCS file resolution from FlowJo URI references |
| R/pretty_print_flowjo.R | Pretty-printing of FlowJo JSON workspace contents for inspection and debugging |
| R/helpers.R, R/helpers-conversion.R | Utility functions, UUID validation, and conversion helpers |

The main exported functions are:

- read_flowjo11_workspace() — parse a FlowJo v11 .flowjo workspace.
- fj11_to_gatingset() — convert a parsed workspace into a list of per-sample GatingSet objects.
- export_flowjo10_workspace() — serialize a GatingSet to FlowJo v10 XML.
- pretty_print_flowjo() — extract and format a .flowjo workspace to human-readable JSON text for debugging.
- set_verbose() / get_verbose() — toggle and query diagnostic output.
