## Supplementary material for "CyFj11: FlowJo v11 Workspace Import and Legacy Format Export for R-Based Flow Cytometry Analysis": Package reference manual, generated from the package's roxygen2 documentation

### Appendix S5: Package Reference Manual

This appendix is the CyFj11 package reference manual generated automatically from the roxygen2 documentation embedded in the R source files. It contains the same content as the rendered man/*.Rd help pages, collected into a single PDF for the manuscript.

#### S5.1 How the reference manual is generated

All exported functions, datasets, and the package overview are documented with roxygen2 comments in the R/ directory. To regenerate the reference manual, run either of the following from the package root:

devtools::document()

or

roxygen2::roxygenize()

This populates the man/ directory with .Rd files. A single PDF can then be produced with:

R CMD Rd2pdf .

The resulting PDF contains one section per help topic and serves as the authoritative function reference for the package. For this manuscript the generated PDF is saved as manuscript/CyFj11_reference_manual.pdf.

#### S5.2 Main topics covered

The reference manual documents the exported functions described in the main text:

- read_flowjo11_workspace() — parse a FlowJo v11 .flowjo workspace
- fj11_to_gatingset() — convert a parsed workspace to a list of GatingSet objects
- export_flowjo10_workspace() — export a GatingSet to FlowJo v10 XML
- pretty_print_flowjo() — pretty-print the JSON content of a .flowjo file
- set_verbose() / get_verbose() — control diagnostic output

It also documents the package purpose, source-file organization, and example data. Technical implementation details (format comparison, gate conversion, transformation mapping) are provided separately in **Appendix S4**.

#### S5.3 File locations

| Generated artifact | Path |
| --- | --- |
| roxygen2 source comments | R/*.R |
| Rendered Rd help pages | man/*.Rd |
| Reference manual PDF | CyFj11.pdf (produced by R CMD Rd2pdf .), saved as manuscript/CyFj11_reference_manual.pdf for the manuscript |
